# Four fundamental processes of community assembly

**DOI:** 10.1101/188607

**Authors:** Joshua Ladau, Steven J. Schwager

## Abstract

A central aim of ecology is understanding the mechanisms of community assembly. To address this problem, community assembly is often modeled as a sampling process, in which species are selected from a pool of available species, possibly with effects of interspecific interactions, habitat filtering, and other ecological mechanisms. However, the fundamental stochastic sampling process by which species are selected from the pool remains unexplored. Here we demonstrate the distinctness of four canonical sampling processes, the Bernoulli, Plackett-Luce, multinomial, and fractional multinomial processes. Each process can be affected by ecological mechanisms or it can occur in their absence. Although all four of the processes are *a priori* plausible and the first two are widely used in ecological models, we show that the multinomial and fractional multinomial processes broadly underlie community assembly.

---

Understanding how communities are structured and assembled constitutes a central aim of community ecology *(1–4)*. Over ecological timescales, community assembly is often considered a sampling process, in which species are selected from a pool of species inhabiting the surrounding areas *(5, 6)*. This process may be affected by dispersal limitation, habitat filtering, interspecific interactions, and other ecological mechanisms. Extensive research has focused on assessing the relative roles of these mechanisms *(7–10)*. However, a fundamental question is which of the many possible general sampling processes govern the selection of species from the pool. Remarkably, this question seems to have received no explicit attention. In this paper, we systematically consider four canonical sampling processes of community assembly with respect to phylogeny, and although all four are reasonable *a priori*, we show that two of them are widely supported by data.

Assessing the fundamental sampling processes of community assembly has important implications: it provides a key baseline for inferring effects of classical ecological mechanisms (e.g., competition and facilitation) from non-experimental data, and it can allow perturbations to communities to be detected and mitigated. Furthermore, an understanding of community assembly processes is important for developing conservation strategies *(11)* and formulating approaches for controlling infectious diseases *(12)*.

Two of the sampling processes that we consider have implicitly been widely used in previous models of community assembly: the Plackett-Luce process and a dependent Bernoulli process *(13, 14)*. They are not equivalent, although this seems not to have been noted previously. The third process, the multinomial, is well known *(15)* but unused in community assembly, and the last process, the fractional multinomial, was described recently *(16)*. These latter two processes are best supported by ecological data.

All four processes posit the existence of a species pool (the set of species that can potentially colonize a community) and phylogenetic information about that pool *(17,18)*. In the first process, the Plackett-Luce *(19)* process, *(i)* each species in the pool has an initial probability of being selected, *(ii)* after the first species is selected, the probabilities of the remaining species are modified to reflect the fact that the first species can no longer be selected, and *(iii)* additional species are subsequently selected, with modifications to the probabilities at each step to reflect that previously selected species are unavailable for selection *(13)*. This process is analogous to drawing marbles (species), possibly with unequal and dependent probabilities, without replacement from an urn (the pool). Effects of habitat filtering are reflected in heterogeneous probabilities (e.g., species A is twice as likely to colonize as species B), while effects of interspecific interactions are reflected in the modifications of the probabilities (e.g., species A is less likely to be drawn if species B is already drawn; an absence of interspecific interactions is modeled by normalizing the remaining probabilities after each draw).

In the second process, the dependent Bernoulli process, each species in the pool has a probability of occurrence, and its outcome (occurs or does not occur) is a Bernoulli random variable *(14)*. This process is analogous to flipping biased coins, potentially non-independently, to determine which species are present. Heterogeneous probabilities again reflect effects of habitat filtering. Dependence among the random variables reflects effects of interspecific interactions; an absence of interspecific interactions is modeled by independence of the random variables.

The third process, the multinomial process, has not been widely applied in community ecology. Let a unit be an arbitrary taxonomic or functional grouping (e.g., genus or family). In the multinomial process, *(i)* species are drawn sequentially and *(ii)* the probability of belonging to each unit in the pool is specified on each draw. Effects of habitat filtering are reflected in the probabilities for the initial draw. Absence of interspecific interactions is modeled by keeping the unit probabilities constant for all draws as in sampling with replacement; while presence of interactions is modeled by modifying the probabilities (e.g., the probability of drawing the second species from unit A might decrease if the first species was drawn from that unit).

The fourth sampling process, until recently not previously considered in community ecology, nor to our knowledge in any other context *(16)*, will now be defined. Together with the multinomial, it explains remarkably well patterns observed in ecological data. Define *〈ij〉* as the event that the *i*th and *j*th species to arrive in the community share the same unit and *〈ij〉*′ as the event that they do not. For a community with three species, the absence of interspecific interactions is modeled by

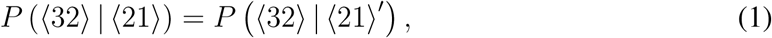

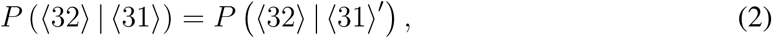

and

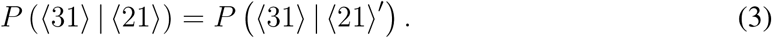

This process is unchanged under effects of habitat filtering, but in the presence of effects of competition or facilitation, each equivalence in (1) - (3) becomes “less than” or “greater than,” respectively. For communities with more than three species, relations analogous to (1) - (3) hold when interspecific interactions are absent, and competition or facilitation have analogous effects on these relations. This process can be shown to be equivalent to a multinomial process in which, by analogy to fractal dimensions, units are fractional. For this reason, we will refer to this process as the fractional multinomial process. Examples of all four processes are given in the Online Supplementary Material.

The four processes are non-equivalent. To demonstrate this, let *I* be a multiset that gives the number of species in each unit in a community; i.e., *I* is an integer partition. For example, if there are four species in a community, two sharing a unit and the third and fourth in separate units, then *I* = {2,1,1} (the other possible partitions of four species are denoted {4}, {3,1},{2, 2},and {1,1,1,1}). Given a species pool, the corresponding set of occurrence probabilities, and the observed number of species, in the absence of interspecific interactions it is straightforward to compute *P(I)* for each possible *I* for the Plackett-Luce, Bernoulli, and multinomial processes. For the fractional multinomial process, it can be shown that (1), (2), and (3) are equivalent to
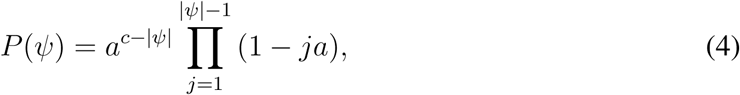
where *c* = 3, *a* ∈ [0,1/2] ∪ {1}, and *ψ* is a set partition of species, as identified by order of arrival, into units *(16)*. Thus, *P*({2,1}) = 3a(1 — a). This result generalizes to arbitrary c ∈ ℕ, where *a* ∈ [0, 1/(c – 1)] ∪ {l/(c – 2)} ∪ … ∪ {1/2} ∪ {1}. Additionally, (4) implies that conditional on c species occurring,

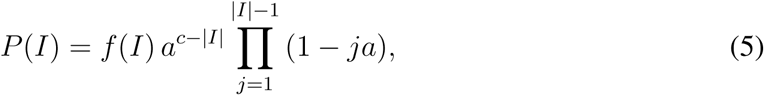

where *f({*i*_*1*_,*i*_*2*_, …,*i*_|*I*|_})* is the number of set partitions with units of sizes *i*_*1*_,*i*_*2*_, …,*i*_|*I*|_ *(16)*. Hence, for any pair of integer partitions *I*_*1*_ and *I*_*2*_, each process defines a function *z*(*θ*) *≡* (*Pθ*(*I*_*1*_), *Pθ*(*I*_*2*_)), where *θ* is a vector of possible parameter values (e.g., occurrence probabilities). Examining the images of these functions reveals that all four of the processes are distinct from one another (Figure 1).

**Figure 1:**
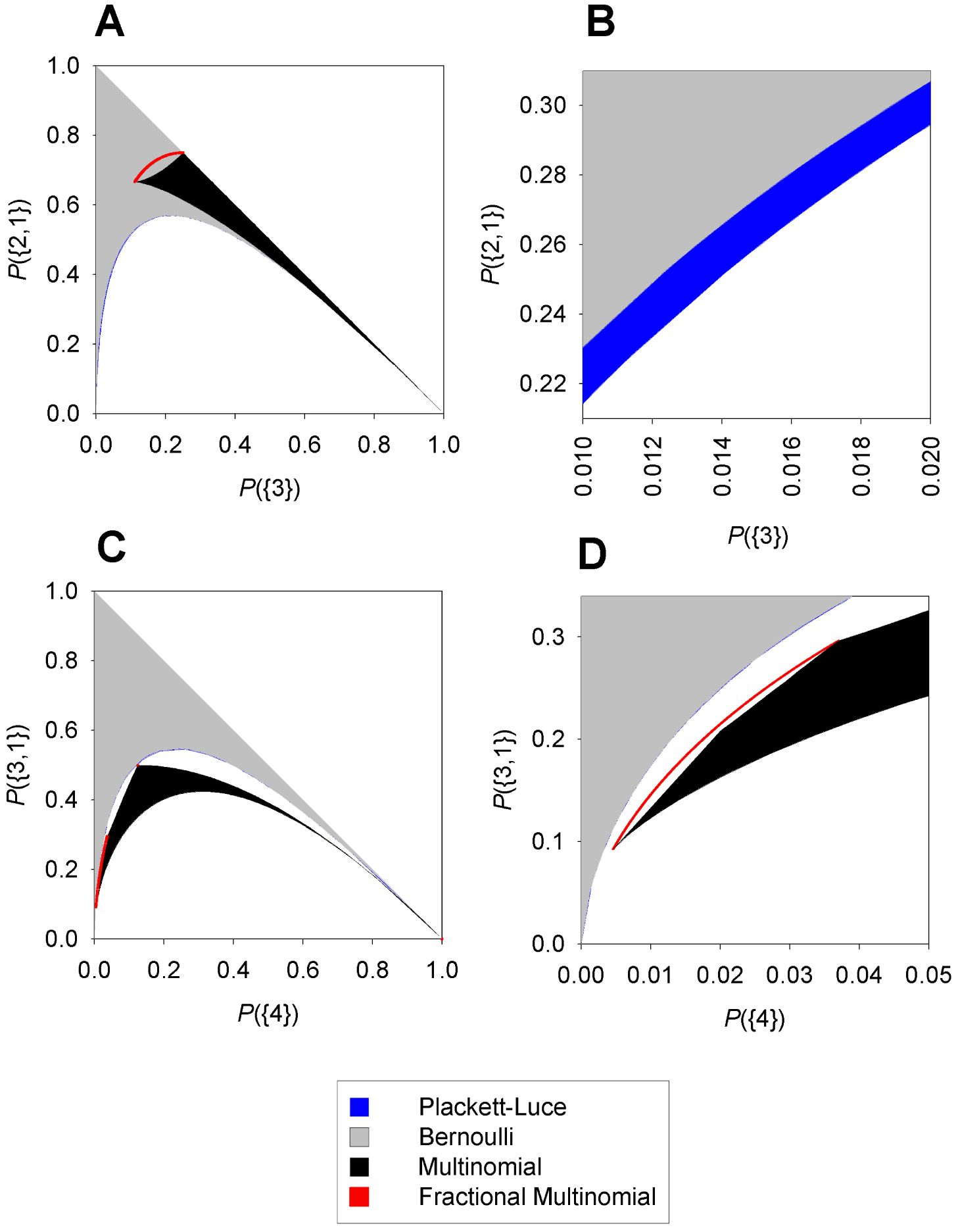
Plots showing that the four sampling processes are distinct. {i, j, k, …} is a multiset giving the observed partition of species into units (e.g., {2, 1} means that 3 species are observed, 2 in the same unit and 1 in a different unit). Probabilities are conditional on the observed number of species (e.g., P({2, 1}) is conditional on 3 species being observed). **(A)** Plot showing that the multinomial, fractional multinomial, and Bernoulli/Plackett-Luce processes are non-equivalent. Sampling is from a pool of 10 species with partition {8, 1, 1}. **(B)** Expanded view of plot A, showing that the Bernoulli and Plackett-Luce processes are distinct. **(C)** Plot for sampling from a pool of 10 species with partition {4, 2, 1, 1, 1, 1}. **(D)** Expanded view of plot C, showing the that the multinomial, fractional multinomial, and Bernoulli/Plackett-Luce processes are distinct. See Online Supplementary Materials for additional details.

*A priori,* theoretical considerations do not indicate which of the processes should occur in ecological communities; assembly could follow any of the possible patterns specified by these four models. To address this question, we investigated the consistency of the four processes with two large data sets: the Barro Colorado Island 50Ha vegetation census (“BCI”) *(20-22)* and the North American Breeding Bird Survey (“BBS”) (*23*). For the BCI data, we divided the 50Ha plot into 1.75m quadrats and treated each quadrat as a site. For the BBS data, we reasoned that assembly processes may differ regionally, and that they should be most distinguishable in communities that are close to equilibrium; i.e., communities of non-migratory species. Hence, we considered separately the data from two representative bird conservation regions, and we performed analyses for permanent resident species. We used data only from highly-rated routes, treated each stop along each route as a site, and employed taxonomic information from the most recent AOU checklist. For all data sets, we performed analyses at three taxonomic resolutions: order, family, and genus. For each data set and at each taxonomic resolution, we compared the empirical distributions of I with fitted distributions derived from each sampling process (see Online Supplementary Material).

In all cases, except when units were defined as orders for the BBS data, the multinomial and fractional multinomial processes were most consistent with the data. They accounted for a remarkable proportion of variation in the observed frequencies of *I* (median *R*^*2*^ > 0.999; minimum *R*^*2*^ = 0.98), were strongly favored over the Plackett-Luce and Bernoulli processes in an AIC analysis *(24)*, and unlike these other two processes, were never rejected in likelihood ratio goodness-of-fit tests (Table 1). The fractional multinomial process was favored over the multinomial process in two out of seven cases by a likelihood ratio test; in the other cases, there was insufficient power to distinguish between the processes. When units were defined as orders for the BBS data, all models were inappropriate (*R*^*2*^ ≈ 0 in all cases, *p* < 0.0001 in goodness of fit tests). This demonstrated that the aforementioned close fits were not a trivial artifact of the fitting procedure.

**Table 1:** Goodness-of-fit for the sampling processes. BCI and BBS refer to the 50Ha Barro Colorado Island Vegetation census and the North American Breeding Bird Survey, respectively. BBS 5 and 16 refer to bird conservation regions 5 and 16. Units were defined as families for the results presented here; see Online Supplementary Material for the results using genera and orders. Marginal *R*^*2*^ is the proportion of variation left unexplained by the Plackett-Luce/Bernoulli processes that the Multinomial and Fractional Multinomial processes explained. *P* is the *p*-value from a likelihood ratio goodness-of-fit test. Some of the reported values are bounds or approximations; see Online Supplementary Material for details. Overall, the data are most consistent with the multinomial and fractional multinomial processes.

| Data | Process | AIC | $R^2$ | Marginal $R^2$ | $P$ |
| --- | --- | --- | --- | --- | --- |
| BCI | Plackett-Luce/Bernoulli | $\geq 50252.11$ | $\geq 0.99$ | 0 | 1 |
| | Multinomial | $\leq 49696.32$ | $\geq 0.99$ | 0.97 | 1 |
|  | Fractional Multinomial | 49696.32 | 0.99 | 0.97 | 1 |
| BBS 5 | Plackett-Luce/Bernoulli | $\geq 4091.09$ | $\geq 0.91$ | 0 | $\geq 0$ |
| | Multinomial | $\leq 4024.32$ | $\geq 0.99$ | 0.99 | $\geq 0.99$ |
|  | Fractional Multinomial | 4023.99 | 0.99 | 0.99 | 0.99 |
| BBS 16 | Plackett-Luce/Bernoulli | $\geq 4661.93$ | $\geq 0.34$ | 0 | $\geq 0$ |
| | Multinomial | $\leq 4622.23$ | $\geq 0.98$ | 0.97 | $\geq 0.13$ |
|  | Fractional Multinomial | 4606.09 | 0.99 | 0.99 | 0.13 |

Numerous studies have examined whether communities are consistent with “random” sampling from species pools, with the intent to use deviations from “randomness” as a means for assessing the effects of interspecific interactions, habitat filtering, and other ecological processes **(8, 17, 18)**. Unlike these studies, the present study evaluated which of four possible fundamental sampling processes governed community assembly. These sampling processes can occur in both the presence and absence of the aforementioned ecological phenomena, and they provide an overarching framework for modeling community assembly as a selection process from species pools.

The analyses required three assumptions: first, that data from different sites were independent; second, that data were identically distributed; and third, that interspecific interactions did not affect the assembly processes. All of these assumptions were surely violated to some extent *(25, 26)*, but the extremely close fits for the multinomial and fractional multinomial processes suggest that these violations were negligible; we were able to clearly show that these two processes governed community assembly from among the other two possibilities.

The analyses indicate that the fractional multinomial process, and potentially the multinomial process, underlie community assembly very broadly. The data sets that we examined were at disparate spatial scales (10^1^ to 10^3^m) and for disparate taxa (plants and birds), yet they were both highly consistent with the two processes. Analyses of other data sets, not presented here, bolster this conclusion. Differing results are likely to provide important insight into ecological phenomena, environmental heterogeneity, and new sampling processes.

Why might the multinomial and fractional multinomial process be so prevalent? The fractional multinomial process specifies only general independence properties for the sampling process [equations (1) to (3)], unlike the other processes, which posit different and more narrowly specified properties. Although all of the processes are plausible, the unique generality of the fractional multinomial process may account for its prevalence.

Sampling processes are of great practical importance. Most existing models of community assembly assume either the Plackett-Luce or Bernoulli process, and these are used as if they were interchangeable *(13,14)*. However, the nature of the underlying process can have profound implications: it can constrain the ecological phenomena that can be observed, as well as the inferences and predictions that can be made. By explicitly considering sampling processes, it will be possible to make more effective inferences in ecology *(16)*.

Most research on community assembly has focused on specific “assembly rules” and the influence of ecological mechanisms on the assembly process (1,7,17,18). Until now, the sampling process by which communities assemble has received no explicit attention; rather, in modeling community assembly, sampling processes have been implicitly assumed. Our results indicate that assembly occurs via two heretofore unconsidered processes: the multinomial and fractional multinomial processes. Further research on these sampling processes will elucidate additional fundamental mechanisms of community assembly.

## Online Supplementary Material

### Example: Computing Distributions

This section presents an example of how to compute the distributions predicted by the four sampling processes. A species pool of five species is assumed in the example. Species 1, 2, and 3 are in genus *A*, and species 4 and 5 are in genus *B*. A lack of interspecific interactions is also assumed; that species are drawn independently. The Plackett-Luce process posits that species are drawn sequentially, and that each species in the pool has a specified probability of being selected on the first draw. Denoting the probability for species *i* by *p*_*i*_, we have the constraint that *p*_*1*_ + *P*_*2*_ + *P*_*3*_ + *P*_*4*_ + *P*_*5*_ = 1 (one of the five species must be picked on the first draw). The probability of observing the partition {2, 1} is given by

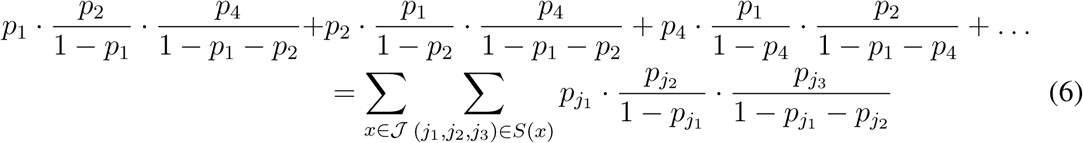

where

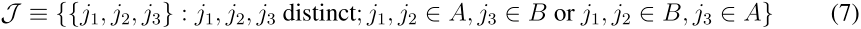

and for any set *E*, *S(E)*is the set of all permutations of *E*. The Bernoulli process posits that species *i* has a probability of occurring here denoted *q*_*i*_. Hence, 0 ≤ *q*_*i*_ ≤ 1, and the probability of {2, 1} is

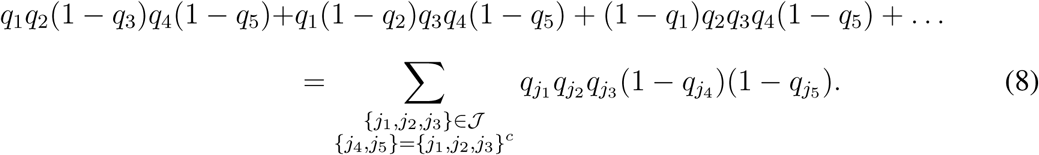

To compare the binomial process to the other processes, it is necessary to compute the probability of {2,1} conditional on observing 3 species; i.e., dividing *(8)* by the probability of observing 3 species. The multinomial process posits that species are drawn sequentially, and that each species has a specified probability of belonging to each genus in the pool. Letting *r*_*j*_ denote the probabilities associated with genus *j*, we have the constraint *r*_*A*_ + *r*_*B*_ = 1 and the probability of {2, 1} is

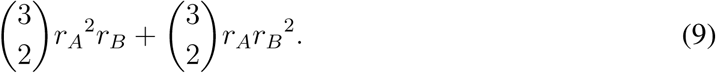

The fractional multinomial has a single parameter *a*, and regardless of the composition of the pool, P({2, 1}) = 3a(1 — a), with *a* ∈ [0,1/2] ∪ {1}.

### Process Non-equivalence

This section details how we plotted the images of *z*{*θ*) ≡ (*Pθ*(*I*_*1*_), *Pθ*(*I*_*2*_)) for each process, where *θ* is a vector of parameter values, and *I*_*1*_ and *I*_*2*_ are integer partitions. The plots of these images are shown in Figure 1. For the fractional multinomial process, the images are parametrically defined curves, which can be plotted routinely. For the other processes, our strategy was to find boundary points, from which we could easily interpolate the shapes of the images. For a given process and vector of parameter values *θ* (e.g., probabilities of species occurring in the binomial process), let *P*_*θ*_(*I*) give the probability of integer partition *I*, conditional on *c* species occurring. Let Θ be the set of all possible parameter values; i.e., *θ* ∈ Θ. For any point (*x*_*o*_,*y*_*o*_) ∈ [0, l]^2^ and integer partitions *I*_*1*_ and *I*_*2*_, if *θ* ∈ Θ satisfies the following two conditions, then *(*Pθ*(*I*_*1*_)*, *Pθ(*I*_*2*_))* is a boundary point:

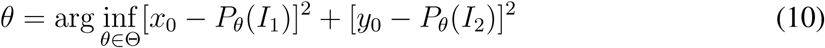

and

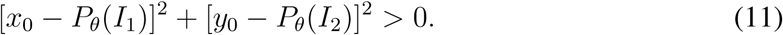

If 5 or fewer species are sampled from a pool of fewer than 11 species, closed form expressions can be found for *Pθ(I)* on Θ. These are generally polynomials and quotients of polynomials with several hundred to several thousand terms. We were able to programmatically generate string expressions for these polynomials, and then minimize *[*x*_*0*_ — *Pθ*(*I*_*1*_)]^2^* + *[*j*_*0*_ — *Pθ*(*I*_*2*_)]^2^* for values of *(*x*_*0*_,*y*_*0*_)* using simulated annealing in Mathematica. We accepted *(*Pθ*(*I*_*1*_, *Pθ*(*I*_*2*_))* as a boundary point only if *[*x*_*0*_ — *Pθ*(*I*_*1*_)]^2^* + *[*y*_*0*_ — *Pθ*(*I*_*2*_)]^2^* > 0.01. We considered values of *(*x*_*0*_,*y*_*0*_)* along 0.01 increment (multinomial process) and 0.05 increment grids (Bernoulli and Plackett-Luce processes). Code was written in Visual Basic 6.0 and Mathematica 6.0, and is available from J.L.

## Data Analysis

The primary aim of our data analyses was to assess which sampling process (Plackett-Luce, Bernoulli, multinomial, or fractional multinomial) was most consistent with the data. We used four measures for making this assessment: The first measure was *R*^*2*^, the proportion of variation in the observed frequencies of / attributable to the process. Following **(27)**, we defined *R*^*2*^ as

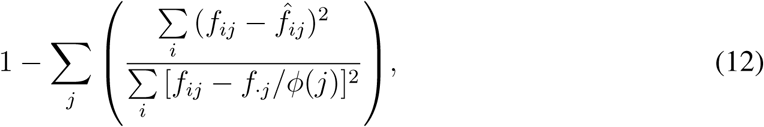

where *f*_*ij*_ is the observed frequency of partition *i* given the presence of *j* species, 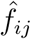 is the frequency predicted by the maximum likelihood fit of the model, *f*._*j*_ is the total number of partitions observed with j species, and *Φ(*j*)* gives the number of partitions of *j* species. We used *R*^*2*^ as a general measure of goodness of fit: if *R*^*2*^ was less than 0.5, we took this as evidence that assumptions of site independence and similarity were violated or that the sampling process was inappropriate. The second measure that we used was a likelihood ratio goodness-of-fit test. A significant result from this test also provided evidence that assumptions were violated or the sampling process was inappropriate. We primarily employed the third and fourth measures to distinguish which of the four processes was *most* appropriate. When the processes had different numbers of parameters, we used AIC; when they had the same number of parameters, we used a likelihood ratio test.

All of these measures require maximizing the likelihood function for each process. In our analyses, we assumed that the observations at different sites were independent and identically distributed. (As discussed in the text, the analyses were robust to this assumption.) Under this assumption, the Plackett-Luce and Bernoulli model have as many parameters as there are species in the pool (≈ 100 to 1000). The multinomial process has as many parameters as there are units in the pool (≈ 25), and the fractional multinomial has one parameter.

Because of the complexity of the first three models, we were unable to maximize the likelihood functions reliably, despite extensive efforts. Hence, we pursued two alternative approaches. The first approach was to find an upper bound for the likelihood functions. The upper bound that we used is given by:

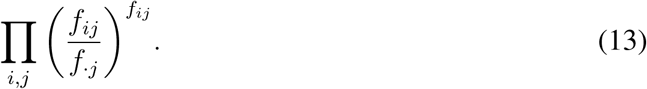

This bound is useful for comparing AIC values, but not for computing values of *R*^*2*^ or performing goodness-of-fit tests. Specifically, for the AIC, if a model has *κ* parameters and the upper bound for the likelihood function is λ*, then a lower bound on the AIC value is —2log(λ*) + *2κ*.

The second approach was to find maximum likelihood estimates for simpler versions of the processes. The simpler versions that we considered were:

1. The Plackett-Luce process, assuming that all species are equally likely to be selected. The numbers of species in the units in the pool were free parameters.
2. The Bernoulli process, assuming that all species are equally likely to occur. As above, the numbers of species in the units in the pool were free parameters.
3. The multinomial process, assuming that species are equally likely to be drawn from each unit. The number of units is a free parameter.

For these simpler models, we were able to calculate *R*^*2*^ and *p*-values from likelihood ratio goodness of fit tests. We also calculated upper bounds on AIC. If a simplified model fit extremely poorly (e.g., *R*^*2*^ ≈ 0 and *p* ≈ 0), we judged that the full model was likely to fit poorly too.

## Detailed Results

Tables 2 and 3 list the results when units were defined as genera and orders; Table 1 (printed text) gives the results for families. For the Barro Colorado Island data (n = 85059), all of the simplified processes fit well, having high values of *R*^*2*^ and never being rejected in goodness-of-fit tests. However, the full multinomial and fractional multinomial processes were clearly favored over the full Plackett-Luce and Bernoulli processes: the upper bound for the AIC value of the full multinomial model was less than the lower bound for the Plackett-Luce and Bernoulli processes, and the exact AIC value for the fractional multinomial was less than the lower bound for the Plackett-Luce and Bernoulli processes. Despite having only one parameter, the fractional multinomial always had the highest *R*^*2*^ values. For the Breeding Bird Survey data (*n = 4249* and *3707* for bird conservation regions 5 and 16, respectively), the simplified Plackett-Luce and Bernoulli processes were always rejected in goodness-of-fit tests. When units were defined as orders, the multinomial and fractional multinomial processes were also rejected, and all processes had very low values of *R*^*2*^, indicating that none were appropriate. In the other cases, the full multinomial and fractional multinomial models were again favored by AIC, and the fractional multinomial had the highest values of *R*^*2*^. Overall, the multinomial and fractional multinomial processes were generally favored by the data.

**Table 2:** [Online Supplementary Material] Goodness-of-fit for the sampling processes, with units defined as genera. BCI and BBS refer to the 50Ha Barro Colorado Island Vegetation census and the North American Breeding Bird Survey, respectively. BBS 5 and 16 refer to bird conservation regions 5 and 16. Marginal R^2^ is the proportion of variation left unexplained by the Plackett-Luce/Bernoulli processes that the Multinomial and Fractional Multinomial processes explained. *P* is the *p*-value from a likelihood ratio goodness-of-fit test. Overall, the data are most consistent with the multinomial and fractional multinomial processes.

| Data | Process | AIC | $R^2$ | Marginal $R^2$ | $P$ |
| --- | --- | --- | --- | --- | --- |
| BCI | Plackett-Luce/Bernoulli | $\geq 6902.91$ | $\geq 0.99$ | 0 | 1 |
| | Multinomial | $\leq 6289.07$ | $\geq 0.99$ | 0.99 | 1 |
|  | Fractional Multinomial | 6289.07 | 0.99 | 0.99 | 1 |
| BBS 5 | Plackett-Luce/Bernoulli | $\geq 1713.37$ | $\geq 0.99$ | 0 | $> 0$ |
| | Multinomial | $\leq 1660.88$ | $\geq 0.99$ | 0.13 | $\geq 0.66$ |
|  | Fractional Multinomial | 1660.86 | 0.99 | 0.18 | 0.66 |
| BBS 16 | Plackett-Luce/Bernoulli | $\geq 783.18$ | $\geq 0.99$ | 0 | $\geq 0.01$ |
| | Multinomial | $\leq 693.73$ | $\geq 0.99$ | 0.83 | 1 |
|  | Fractional Multinomial | 693.73 | 0.99 | 0.83 | 0.99 |

**Table 3:** [Online Supplementary Material] Goodness-of-fit for the sampling processes, with units defined as orders. See Table 2 for a full description.

| Data | Process | AIC | $R^2$ | Marginal $R^2$ | $P$ |
| --- | --- | --- | --- | --- | --- |
| BCI | Plackett-Luce/Bernoulli | $\geq 66360.65$ | $\geq 0.99$ | 0 | 1 |
| | Multinomial | $\leq 65821.49$ | $\geq 0.99$ | 0.92 | 1 |
|  | Fractional Multinomial | 65811.92 | 0.99 | 0.97 | 1 |
| BBS 5 | Plackett-Luce/Bernoulli | $\geq 6328.03$ | $\geq 0$ | 0 | $> 0$ |
| | Multinomial | $\leq 9457.77$ | $\geq 0$ | 0.59 | $> 0$ |
| | Fractional Multinomial | 9457.77 | 0 | 0.59 | $< 0.0001$ |
| BBS 16 | Plackett-Luce/Bernoulli | $\geq 3784.01$ | $\geq 0$ | 0 | $> 0$ |
| | Multinomial | $\leq 8280.44$ | $\geq 0$ | 0.6 | $> 0$ |
| | Fractional Multinomial | 8280.44 | 0 | 0.6 | $< 0.0001$ |

This research was funded by a Santa Fe Institute Postdoctoral Fellowship to J. Ladau. The authors thank J. Harte (UC Berkeley) and the researchers at the Santa Fe Institute, particularly J. Wilkins and A. Clauset, for helpful discussion and advice. The BCI forest dynamics research project was made possible by National Science Foundation grants to Stephen P. Hubbell: DEB-0640386, DEB-0425651, DEB-0346488, DEB-0129874, DEB-00753102, DEB-9909347, DEB-9615226, DEB-9615226, DEB-9405933, DEB-9221033, DEB-9100058, DEB-8906869, DEB-8605042, DEB-8206992, DEB-7922197, support from the Center for Tropical Forest Science, the Smithsonian Tropical Research Institute, the John D. and Catherine T. MacArthur Foundation, the Mellon Foundation, the Celera Foundation, and numerous private individuals, and through the hard work of over 100 people from 10 countries over the past two decades. The plot project is part the Center for Tropical Forest Science, a global network of large-scale demographic tree plots.

## References and Notes

1. R.T. Paine, Oecologia 15, 93–120 (1974).

2. M. Tokeshi, Advances in Ecological Research 24, 111–186 (1993).

3. J.A. Drake, The American Naturalist 137, 1–26 (1991).

4. E. Weiher, P. Keddy, Eds., Ecological Assembly Rules: Perspectives, Advances, Retreats (Cambridge University Press, Cambridge, 2001).

5. S.P. Hubbell, The Unified Neutral Theory of Biodiversity and Biogeography (Princeton University Press, Princeton, NJ, 2001).

6. R.H. MacArthur, E.O. Wilson The Theory of Island Biogeography (Princeton University Press, Princeton, NJ, 2001).

7. J.M. Diamond, in Ecology and Evolution of Communities, M.L.Cody, J.M. Diamond, Eds., (Harvard University Press, Cambridge, MA, 1975), pp. 342–444.

8. E.F. Connor, D. Simberloff, Ecology 60, 1132–1140 (1979).

9. N.J. Gotelli, D.J. McCabe, Ecology 83, 2091–2096 (2002).

10. J. Fargione, C.S. Brown, D. Tilman, Proceedings of the National Academy of Sciences of the United States of America 15, 8916–8920 (2003).

11. W. Atmar, B.D. Patterson, Oecologia 96, 373–382 (1993).

12. C. Stephens, paper presented at the Santa Fe Institute, Santa Fe, NM, 6 March 2008. URL precedings.nature.com/documents/1495/version/1/files/npre20081495-1.pdf.

13. N.J. Gotelli, Ecology 81, 2606–2621 (2000).

14. M.E. Gilpin, J.M. Diamond, Oecologia, 75–84 (1982).

15. N.L. Johnson, S. Kotz, N. Balakrishnan, Discrete multivariate distributions (Wiley, New York, NY, 1997).

16. J. Ladau, S.J. Schwager, Journal of Mathematical Biology, 57, 537–555 (2008).

17. C.O. Webb, D.D. Ackerly, M.A. McPeek, M.J. Donoghue, Ann.Rev. Ecol. Syst. 33, 475–505 (2002).

18. B.J. Enquist, J.P. Haskell, B.H. Tiffney, Nature 419, 610–613 (2002).

19. R.L. Plackett, Applied Statistics 24, 193–202 (1975).

20. S.P. Hubbell, R. Condit, R.B. Foster, Barro Colorado Forest Census Plot Data, URL http://ctfs.si/edu/datasets/bci (2005).

21. R. Condit, Tropical Forest Census Plots (Springer-Verlag and R. G. Landes Company, Berlin, Germany, and Georgetown, Texas, 1998).

22. S.P. Hubbell, R.B. Foster, S.T. O’Brien, K.E. Harms, R. Condit, B. Wechsler, S.J. Wright, S. Loo de Lao, Science 283, 554–557 (1999).

23. J.R. Sauer, J.E. Hines, J. Fallon The North American Breeding Bird Survey, Results and Analysis 1966 - 2006, Version 10.13.2007 (2007).

24. K.P. Burnham, D.R. Anderson, Model selection and multimodel inference: A Practical Information-Theoretic Approach (Springer, New York, NY, 2002).

25. J.B. Plotkin, M.D. Potts, N. Leslie, N. Manokoran, J. Lafrankie, P.S. Ashton, Journal of Theoretical Biology 207, 81–99 (2000).

26. K.E. Harms, R. Condit, S.P. Hubbell, R.B. Foster, Journal of Ecology 89, 947–959 (2001).

27. T.O. Kvalseth, The American Statistician 39, 279–285 (1985).

